# High-frequency sampling rate reduces TMS-pulse artifact duration but not decay artifact: implications for immediate TMS-EEG responses

**DOI:** 10.1101/2025.03.05.641655

**Authors:** Antonietta Stango, Agnese Zazio, Guido Barchiesi, Natale Salvatore Bonfiglio, Marta Bortoletto

**Author notes:** Corresponding author: Dr. Marta Bortoletto, PhD, Neurophysiology Lab, IRCCS Istituto Centro San Giovanni di Dio Fatebenefratelli, via Pilastroni 4, 25125 Brescia, Italy.

## Abstract

In studies combining transcranial magnetic stimulation and electroencephalography (TMS-EEG), two artifacts appear instantly after the TMS pulse, i.e., the TMS-pulse Artifact and the Decay Artifact, and limit the possibility to measure immediate cortical excitability responses. High-frequency sampling rates in EEG recordings have shown promise in reducing artifact duration, allowing more rapid signal recovery, which is crucial for developing biomarkers for neuropsychiatric conditions. However, the features of early TMS-induced artifacts for sampling rates above 5000 Hz are still unclear. Here, we explored the duration of TMS artifacts in the first milliseconds after TMS to understand how they can be further reduced in future studies. We recorded from a phantom head model and from a simple electrical circuit with a sampling rate of 4800 Hz, 9600 Hz, and 19200 Hz and at three TMS intensities (40%, 70%, 100% of maximum stimulator output) in two commercial stimulators. Results showed an initial sharp TMS-pulse Artifact lasting less than 1 ms and decreasing in duration at higher sampling rates. However, the signal was back to baseline at about 2-3 ms due to the presence of a decay artifact that was evident even in optimal conditions of low impedance and mostly dependent on stimulation intensity. These results highlight the need to develop efficient ways to eliminate the decay artifact in order to measure immediate TMS responses.

## Introduction

In TMS-EEG coregistration studies, artifacts start and contaminate the EEG signal as soon as the TMS pulse is delivered (Ilmoniemi and Kičić, 2010). The first ones to occur, henceforth “Early Artifacts”, consist of the TMS-pulse artifact and the Decay artifact. They are due to electromagnetic phenomena and are among the most challenging artifacts to remove (Hernandez-Pavon et al., 2023; Rogasch et al., 2017). Understanding how to eliminate or reduce these artifacts and recover the signal occurring immediately after the TMS pulse is crucial for the development of biomarkers of cortical excitability to apply in the study of neuropsychiatric disorders.

The TMS-pulse Artifact is the direct recording of the electromagnetic field generated by the TMS coil in the EEG electrodes (Veniero et al., 2009). This artifact occurs at the time of the pulse, and its duration is mainly determined by the EEG amplifier settings. The amplitude of the artifact is thousands of times greater than the EEG signal and it is not stable enough over trials to be subtracted, even when inter-trial variability is reduced by synchronizing TMS pulse delivery and EEG sampling (Jamil et al., 2024; Tomasevic et al., 2017). Consequently, at present, there is no reliable method to subtract the TMS pulse artifact from the signal, which means that the signal immediately following the TMS is lost (Hernandez-Pavon et al., 2023).

The Decay Artifact starts as soon as the TMS pulse is delivered and slowly decreases with a power law function (Freche et al., 2018). It is thought to derive from the polarization of the gel and the electrodes over the skin (Litvak et al., 2007). Its duration is positively related to the electrode-skin impedance and stimulation intensity (Freche et al., 2018; Veniero et al., 2009). For high-impedance recordings and for active electrode systems it can reach up to hundreds of milliseconds, whereas an accurate cleaning of the skin can reduce it to a few milliseconds. Various offline processing methods have been developed to minimize this artifact (Casula et al., 2017; Freche et al., 2018; Litvak et al., 2007; Rogasch et al., 2014), but it remains a significant issue in TMS-EEG studies. Indeed, the initial high-amplitude part of the decay artifact is often removed together with the TMS-pulse artifact: Although a few recent studies recovered the EEG signal after 5 ms and focused on early components of TMS-evoked potentials (Bortoletto et al., 2021; Guidali et al., 2023; Zazio et al., 2022), a recent meta-analysis of TMS-EEG literature over the primary motor cortex showed that typically around 15 ms of signal post-TMS pulse is lost (Beck et al., 2024b).

A thorough investigation of these Early Artifacts was performed in 2009 (Veniero et al., 2009) when the duration of TMS-pulse Artifacts was recorded for different stimulation intensities and different sampling rates, up to 5000 Hz, in a passive electrode EEG system: It was demonstrated that the TMS-pulse Artifact duration was reduced with higher sampling rates, reaching 5 ms at 5000 Hz, and was not affected by stimulation intensity. Additionally, it was shown that the Decay Artifact lasted until 15-20 ms in high-impedance conditions and could be sensibly reduced in ideal low-impedance conditions.

In recent years, EEG amplifiers have undergone significant advancements, with some now capable of reaching sampling rates as high as 100000 Hz. A few studies have recorded the effects of TMS pulses at frequencies above 5000 Hz, once again finding that the duration of the recorded pulse decreases with higher sampling rates and varies based on stimulation intensity (Freche et al., 2018; Jamil et al., 2024).

Interestingly, it has been reported that high-speed amplifiers allow the recovery of the signal 2-3 ms after TMS, allowing the measurement of immediate TMS-related responses after primary motor cortex stimulation, which likely represents local cortical excitability (Beck et al., 2024a;Stango et al., 2024 preprint). This finding holds great potential for the development of new, low-cost biomarkers for neuropsychiatric disorders that involve motor alterations, such as depression. Therefore, a more systematic investigation is necessary to understand the constituting features of these Early Artifacts and how they can be further reduced.

Here, we explored the features of the Early Artifacts by employing two different stimulators, three stimulation intensities and three high sampling frequencies on a melon (Veniero et al., 2009) and a tester (see Data Recordings).

## Materials and methods

### Transcranial magnetic stimulation (TMS)

TMS was delivered using two different stimulators: Magstim Super Rapid^2^ connected to an Alpha B.I. Coil Range 70mm (Magstim Company, UK) and Magpro X100 connected to a C-B60 coil (MagVenture, Denmark). For both stimulators, three stimulation intensities were used in different blocks: 40%, 70% and 100% of maximal stimulator output. Fifty pulses were delivered for each condition. The coil was centered over the target electrode and positioned so that there was approximately 1 mm distance from the electrode. Its position was kept constant by means of a mechanical support (Magic arm Manfrotto).

### Data recordings

A TMS-compatible electroencephalography (EEG) amplifier (g-HIamp, g.tec medical engineering GmbH, Schiedlberg, Austria) was used to record the signal from a melon and from a tester. The melon was used to reproduce a condition with low impedance, i.e. below 5 kOhm, similar to recordings over the skin. Indeed, Veniero et al 2009 showed similar results between recordings on a melon and on the knee (Veniero et al., 2009). The tester consisted of a circuit of three electrodes connected in a triangular shape with a 5 kOhm resistance in each connection. It was used to measure the Early Artifacts under the known and optimal condition of low impedance. For each stimulation condition, the signal was acquired with three sampling rates in different blocks: 4800 Hz, 9600 Hz and 19200 Hz (Fig 1).

**Figure 1.**
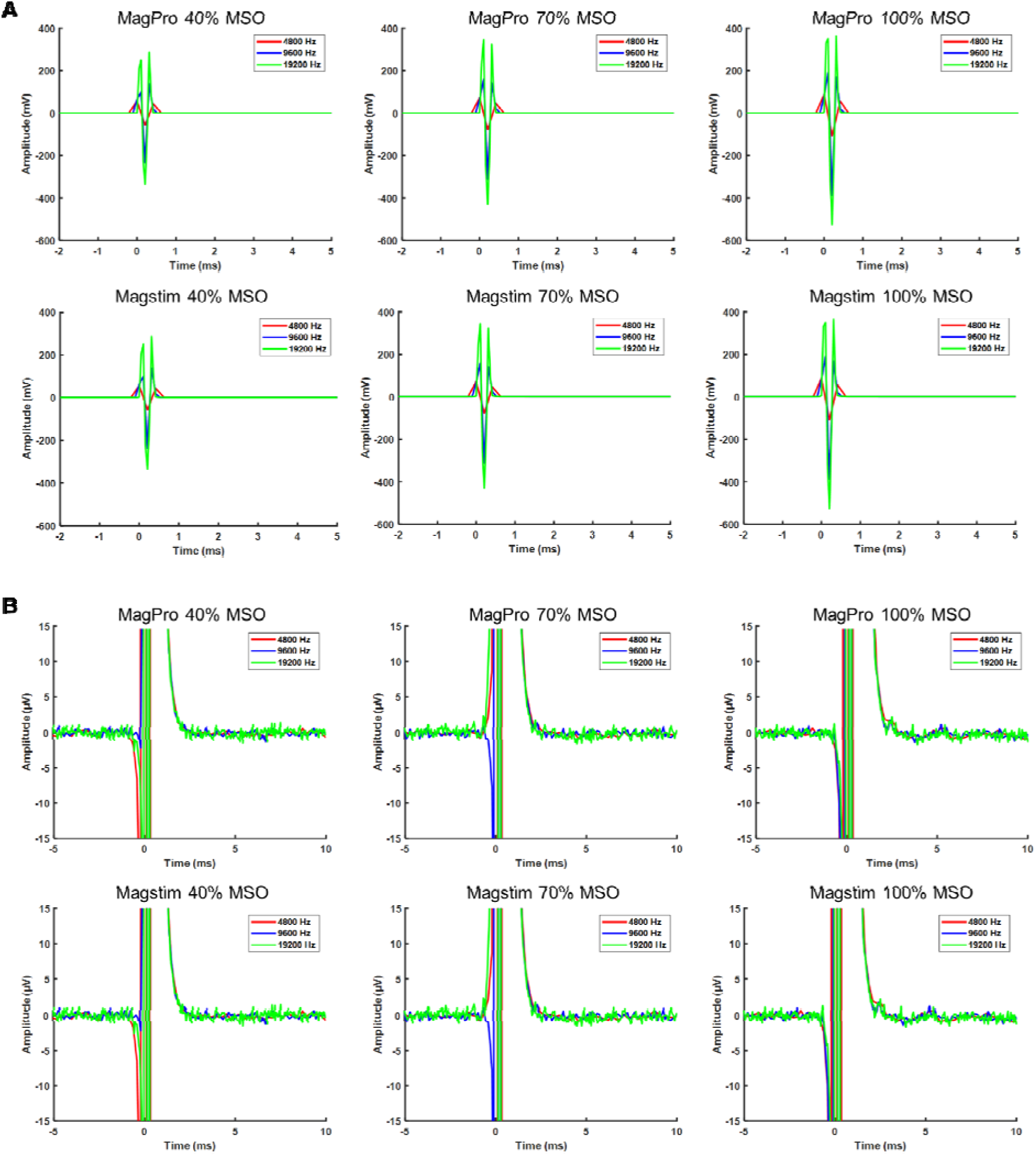
Tester signal averaged over trials in all experimental conditions for MagPro (upper row) and Magstim (lower row). Each plot within a row corresponds to a specific TMS intensity: 40% (left), 70% (middle), and 100% (right) of the maximal stimulator output (MSO). The traces are color-coded to indicate different sampling rates: 4800 Hz (red), 9600 Hz (blue), and 19200 Hz (green). Plots are presented at two distinct axes scales: **A)** the larger scale highlights the TMS-pulse Artifact; **B)** the smaller scale highlights also the Decay Artifact.

### Analysis

Data were preprocessed with Matlab version 2022b (The Mathworks, Natick, MA, USA) adopting functions from EEGLAB version 2023.0 (Delorme and Makeig, 2004). The preprocessing included epoching around the TMS pulse, from −10 ms to 20 ms, and applying baseline correction. Moreover, we corrected for a delay in the g.tec amplifier, known as the Analog Signal Line Delay, between the analog biosignal input, e.g. the EEG signal, and digital trigger input line, i.e. the marker of the TMS pulse. This delay causes each marker to anticipate the EEG signal by a fixed number of sampling points, depending on the sampling rate. Specifically, it corresponds to 6 samples at a sampling rate of 4800 Hz, to 11 samples at 9600 Hz, and to 21 samples at 19200 Hz. In the following analysis, we corrected for the Analog Signal Line Delay by subtracting the corresponding number of samples.

To describe the features of each of the Early Artifacts, we proceeded as follows: First, we calculated the total duration of the Early Artifacts from the marker of the TMS pulse, identifying the time point where the signal returns to baseline values (i.e., Back to Baseline - B2B). Second, we calculated the duration of the TMS-pulse Artifact (i.e., End of TMS-pulse Artifact - ETA). In this way, we aimed at separating the temporal contribution of the two components of the Early Artifacts.

To measure the B2B value, we adapted an algorithm described by Jamil et al. (2024). This algorithm identifies the onset and offset of the Early Artifacts by measuring the peak-to-peak amplitude of the signal in a moving window and comparing it to a threshold based on the pre-TMS signal. We set a 1-ms long window moving in steps of 1 sample and a threshold equal to 3 times the interquartile range of the prestimulus signal, as in Jamil et al. (Jamil et al., 2024). The onset time was defined as the midpoint of the first window in which the peak-to-peak value was above the threshold. Similarly, the offset time was marked as the midpoint of the first window after the onset time in which the peak-to-peak value fell below the threshold.

To measure the ETA, we exploited the local slope of the signal (Leach, 2014), calculated as the ratio between the numerical gradient of the signal and the numerical gradient of time (Fig 2). We considered the signal between the onset and the offset of the Early Artifacts as obtained with the B2B method. We expected high absolute local slope values for the TMS-pulse artifact, due to its extremely rapid and high-energy nature, and lower absolute local slope values for the Decay Artifact, with different orders of magnitude between the two artifacts. Moreover, considering that the current waveform of the TMS pulse is biphasic, we expected the absolute local slope values to be nearly symmetrical around the TMS-pulse Artifact peak, i.e., in the same order of magnitude. Therefore, we defined the end of the TMS-pulse Artifact based on a threshold identified from the local slope at the beginning of the artifact. Specifically, this threshold was the absolute slope value measured approximately two samples after the B2B onset and just before the major slope change, i.e. 200. We identified the positive and the negative peaks of the local slope and selected the last peak which exceeded the chosen threshold.

**Figure 2.**
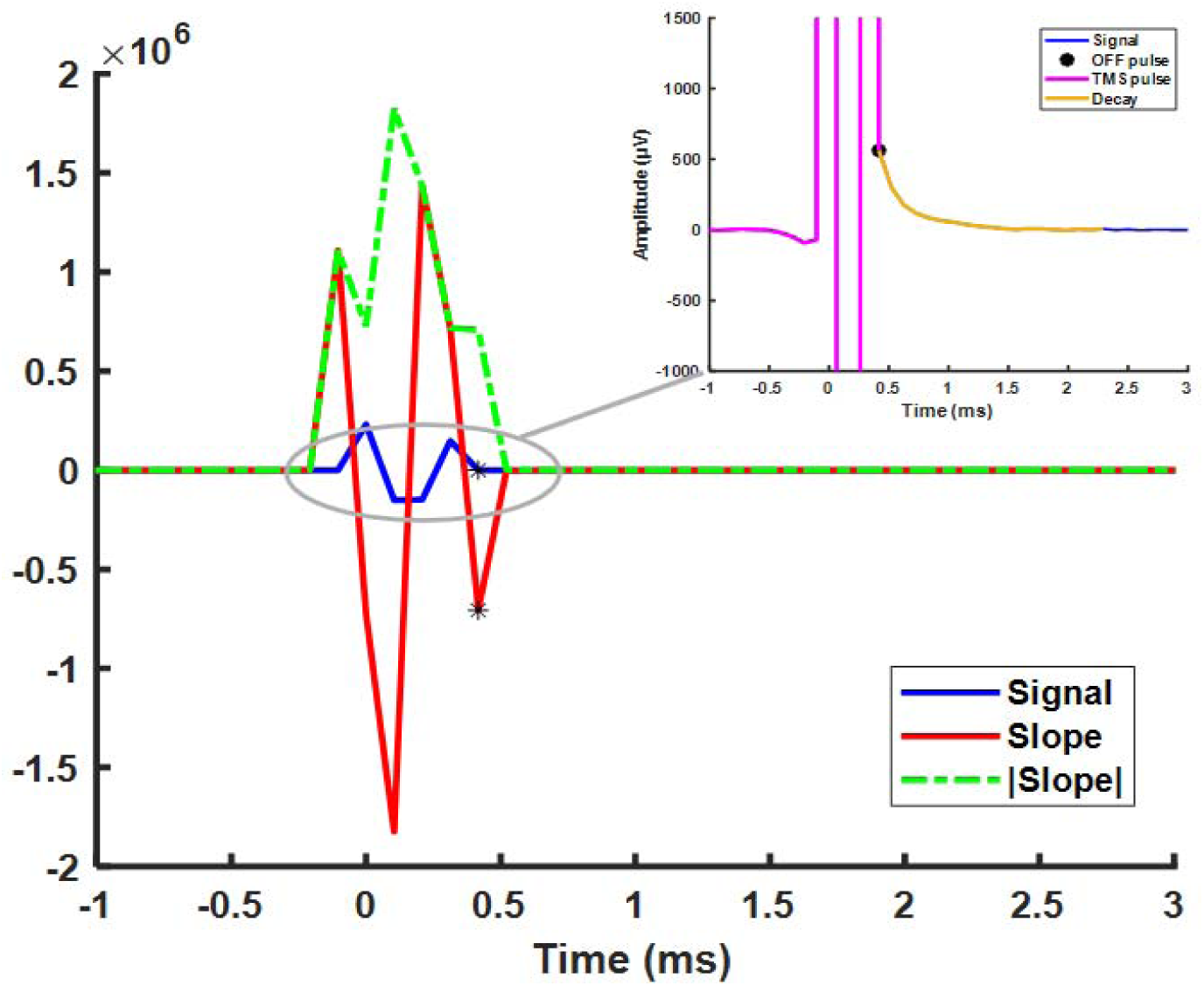
Signal trial (tester- Magpro, Stimulation intensity = 70%, SR=9600) with corresponding slope. The asterisks correspond to the last peak within the threshold for the slope (red line), and to the end of the pulse for the signal (blue line), i.e., TMS-pulse offset. This point marks the beginning of the decay as shown by the black dot in the box on the top right.

### Statistical analysis

As the intensity of the two stimulators are not fully comparable, they were analyzed in separate statistical models. Given that the data distribution deviated from normality (Shapiro-Wilk test, p < 0.05 and further confirmed by a visual inspection of the residual using a Q-Q plots), we applied a non-parametric Aligned Rank Transform Analysis of Variance (ART-ANOVA). ART, as an alternative to the traditional ANOVA applied to rank-transformed data (Elkin et al., 2021; Wobbrock et al., 2011), was used with a 3×3 design, with Sampling rate (4800 Hz, 9600 Hz, and 19200 Hz) and TMS Intensity (40%, 70%, and 100%) as factors, and separately analyzing ETA and B2B as dependent variables. In case of significant interaction, only the simple effects were interpreted, adjusting for multiple testing using Tukey’s correction. The significance level was set at *p* < 0.05.

The R package was used for all statistical analyses (version 4.4.1; https://www.r-project.org/) with the RStudio interface (http://www.rstudio.com/), and ART-ANOVA was implemented with the ARTool package (https://cran.r-project.org/web/packages/ARTool/vignettes/art-contrasts.html).

## Results

We observed different values and different patterns of effects when considering the B2B and ETA values suggesting that the Early artifacts observed in this study included both the TMS-Pulse Artifact and the Decay Artifact. This was also evident looking at the local slope: while the absolute slope values were nearly symmetrical up to a certain point in the final part of the artifact, they eventually leveled off, maintaining the same order of magnitude, indicating that the signal trend had become almost parallel to the x-axis. This trend supports that the nearly symmetrical portion measured with ETA represented the TMS-Pulse Artifact, while the latter portion was the Decay Artifact, the duration of which was measured with B2B.

### Early TMS artifacts duration: Back to baseline (B2B)

The duration of the artifact as B2B value was overall mostly influenced by TMS intensity, with longer durations for higher intensities. Significant effects are illustrated in Figure 3A-B.

**Figure 3.**
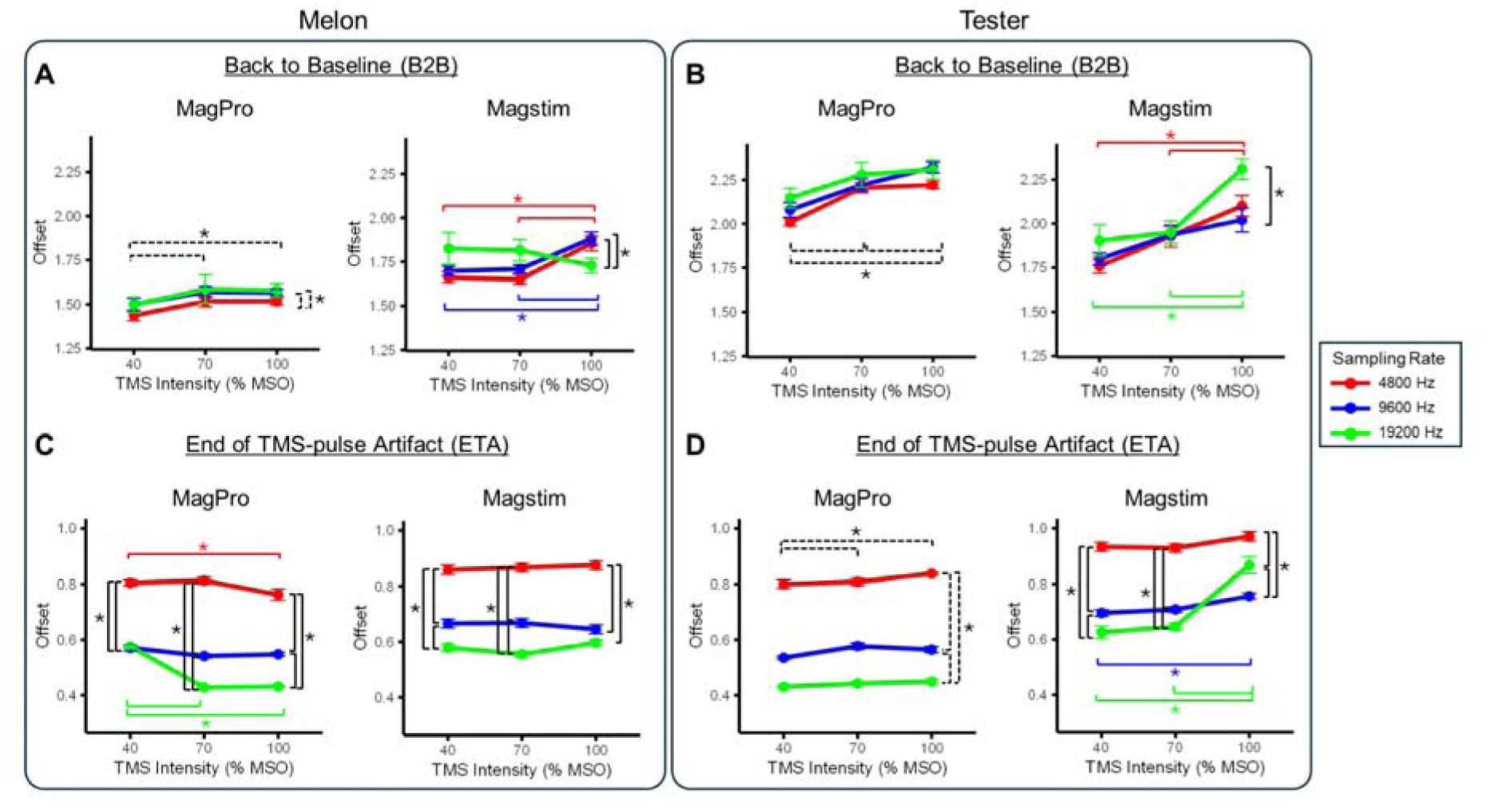
Mean values of artifact duration in the different experimental conditions, from the melon (left panels, A-C) and from the tester (right panels, B-D). Results on the Back to Baseline (B2B) measure are represented in the upper row (**A-B**), and the results of the End of Sharp Artifact (ESA) measure are represented in the lower row (**C-D**). Error bars represent the standard error. Square brackets and asterisks indicate significant effects resulting from main effects (dashed lines) or interaction (solid line).

Specifically, when considering measurements on the melon, the MagPro showed a very similar pattern for all sampling rates with respect to TMS intensity, consisting of a modest but significant increase in artifact duration, ranging on average from 1.48 ms at 40% MSO to 1.55 ms at 100% MSO (main effect of TMS Intensity; *F*_(2,441)_=20.8, *p*<0.001). Similarly, the Magstim showed an increase in artifact duration with TMS intensity for both 4800 Hz and 9600 Hz - but not 19200 Hz - ranging from 1.60 ms at 40% MSO to 1.69 at 100 % MSO (significant TMS Intensity by Sampling rate interaction: *F*_(4,441)_=4.7, *p*=0.001). Overall the Early Artifacts duration was not consistently reduced by sampling rate in either MagPro or Magstim; only in Magstim at 100% MSO artifact duration was significantly shorter in 19200 Hz compared to 4800 Hz (*p*=0.012; *d*=-1.20) and 9600 Hz (*p*<0.001; *d*=-1.12). Recordings on the tester showed a similar pattern, namely an increase in artifact duration with TMS intensity (MagPro: main effect of TMS Intensity, *F*_(2,441)_=42.6, *p*<0.001; Magstim: TMS Intensity by Sampling rate interaction, *F*_(4,436)_=4.67, *p*=0.001, explained by an increase in duration in 4800 Hz and 9600 Hz). Mean B2B values ± standard error are reported in Table 1 and a comprehensive report of the results are available in Tables S1.

**Table 1.** Artifact duration as Back to Baseline (B2B) value. Reported values represent the mean ± standard error over trials for each condition and are expressed in ms.

|  | Sampling rate |  |  |  |  |  |
| --- | --- | --- | --- | --- | --- | --- |
| MELON | MagPro |  |  | Magstim |  |  |
| TMS intensity | 4800 Hz | 9600 Hz | 19200 Hz | 4800 Hz | 9600 Hz | 19200 Hz |
| 40 % | 1.44 ± 0.03 | 1.50 ± 0.03 | 1.50 ± 0.04 | 1.66 ± 0.03 | 1.70 ± 0.03 | 1.82 ± 0.09 |
| 70 % | 1.52 ± 0.03 | 1.57 ± 0.03 | 1.59 ± 0.08 | 1.65 ± 0.03 | 1.71 ± 0.02 | 1.81 ± 0.06 |
| 100 % | 1.52 ± 0.02 | 1.57 ± 0.02 | 1.58 ± 0.04 | 1.85 ± 0.04 | 1.88 ± 0.04 | 1.73 ± 0.04 |
| TESTER | MagPro |  |  | Magstim |  |  |
| TMS intensity | 4800 Hz | 9600 Hz | 19200 Hz | 4800 Hz | 9600 Hz | 19200 Hz |
| 40 % | 2.01 ± 0.02 | 2.08 ± 0.04 | 2.15 ± 0.06 | 1.76 ± 0.04 | 1.80 ± 0.03 | 1.90 ± 0.09 |
| 70 % | 2.20 ± 0.03 | 2.22 ± 0.04 | 2.28 ± 0.07 | 1.93 ± 0.06 | 1.94 ± 0.05 | 1.95 ± 0.06 |
| 100 % | 2.22 ± 0.02 | 2.32 ± 0.03 | 2.31 ± 0.05 | 2.10 ± 0.06 | 2.02 ± 0.07 | 2.31 ± 0.06 |

### End of TMS-pulse Artifact (ETA)

When measured as ETA, the intertrial variability was very low, so that in some conditions the same value was repeated in every trial with a few exceptions, suggesting that the applied method provided a measure with very little measurement error (see Table 2). Overall, the duration of the TMS-pulse Artifact decreased with increasing sampling rates, with a limited impact of TMS Intensity, both in the recordings on the melon and on the tester. Significant effects are illustrated in Figure 3C-D.

**Table 2.** Artifact duration as End of TMS-pulse Artifact (ETA) value. Reported values represent the mean ± standard error over trials for each condition and are expressed in ms.

|  | Sampling rate |  |  |  |  |  |
| --- | --- | --- | --- | --- | --- | --- |
| MELON | MagPro |  |  | Magstim |  |  |
| TMS intensity | 4800 Hz | 9600 Hz | 19200 Hz | 4800 Hz | 9600 Hz | 19200 Hz |
| 40 % | 0.80 ± 0.01 | 0.57 ± 0.01 | 0.58 ± 0.00 | 0.86 ± 0.02 | 0.67 ± 0.02 | 0.58 ± 0.01 |
| 70 % | 0.81 ± 0.01 | 0.54 ± 0.01 | 0.43 ± 0.00 | 0.87 ± 0.02 | 0.67 ± 0.02 | 0.56 ± 0.01 |
| 100 % | 0.76 ± 0.02 | 0.55 ± 0.01 | 0.43 ± 0.00 | 0.87 ± 0.02 | 0.65 ± 0.02 | 0.60 ± 0.01 |
| TESTER | MagPro |  |  | Magstim |  |  |
| TMS intensity | 4800 Hz | 9600 Hz | 19200 Hz | 4800 Hz | 9600 Hz | 19200 Hz |
| 40 % | 0.80 ± 0.02 | 0.54 ± 0.01 | 0.43 ± 0.01 | 0.93 ± 0.02 | 0.70 ± 0.01 | 0.63 ± 0.02 |
| 70 % | 0.81 ± 0.01 | 0.58 ± 0.01 | 0.44 ± 0.01 | 0.93 ± 0.01 | 0.71 ± 0.01 | 0.65 ± 0.01 |
| 100 % | 0.84 ± 0.00 | 0.56 ± 0.01 | 0.45 ± 0.01 | 0.97 ± 0.02 | 0.76 ± 0.01 | 0.87 ± 0.03 |

Specifically, in the melon measurements, artifact duration was significantly shorter with increasing sampling rates for almost all TMS intensities, with the only exception of 9600 Hz vs. 19200 Hz which did not differ in 40% MSO for MagPro and in 100% MSO for Magstim (TMS intensity by Sampling Rate interaction; MagPro: *F*_(4,441)_=43.6, *p*<0.001; Magstim: *F*_(4,441)_=6.03, p<0.001). On average, the reduction of the TMS-pulse Artifact duration with TMS intensity ranged from 0.83 ms at 4800 Hz to 0.53 ms at 19200 Hz. Similarly, results on the tester showed shorter artifact duration for increasing sampling rates for all TMS intensities, except for 100% MSO in Magstim (MagPro: main effect of Sampling Rate, *F*_(4,441)_=885.8, *p*<0.001; Magstim: Sampling Rate by TMS Intensity interaction, *F*_(4,436)_=13.1, *p*<0.001). In the tester, a small but consistent effect of TMS Intensity was also present, with longer TMS-pulse Artifact durations with increasing TMS Intensity (MagPro: main effect of TMS Intensity, *F*_(2,441)_=6.1, *p*=0.002). Mean ETA values ± standard error are reported in Table 2 and a comprehensive report of the results are available in Table S2.

## Discussion

In the present experiment we tested whether and how the sampling rate and the intensity of stimulation affected the duration of Early Artifacts after TMS pulse delivery on the EEG signal. Our data show that the EEG signal immediately after the delivery of a TMS pulse is affected by a very short high-amplitude TMS-pulse Artifact followed by a Decay Artifact of a few milliseconds, so that the signal can be recovered after approximately 2-3 ms, which is a great amelioration compared to only a few years ago. However, the critical point highlighted by our data is that the advantages of the sampling rate increase, and thus of a time-shortened TMS-pulse Artifact, are strongly reduced by the presence of a concomitant and longer-lasting Decay Artifact, virtually not affected by the sampling rate increase.

These results highlight the advancement in reducing the TMS-pulse Artifact by current high-sampling rate amplifiers compared to the state of the art of a few years ago (Veniero et al., 2009). Indeed, the TMS-pulse Artifact duration is affected by the amplifier settings. A crucial role is played by the sampling rate, as shown by our results and by previous studies (Freche et al., 2018; Veniero et al., 2009). The sampling rate is directly linked to the hardware anti-aliasing low-pass filter that reduces the amplitude of the artifact but increases its duration. Therefore, increasing the sampling rate allows to reduce the duration of the artifact until the possibility to measure its actual shape. Regarding this, it should be noted that the anti-aliasing filter associated with sampling rate may change in different amplifiers. For example, the highest low-pass filter in the BrainAmp at 5000 Hz is 1000 Hz and allows recording with a TMS-pulse Artifact of 5 ms. At similar sampling frequency, i.e., 4800 Hz, the g.HIamp has a higher anti-aliasing filter and allows recording a shorter TMS-pulse Artifact. Therefore, commercially available amplifiers may perform differently even at comparable sampling frequencies.

Notably, the highest sampling rate used in this study was not sufficient to measure the TMS-pulse Artifact with its actual shape, i.e. a biphasic curve lasting 300 microseconds, and without distortion. The effect of hardware filters was evident in two features of the artifact: first, its duration was longer than the actual pulse. Second, after applying the correction for the delay of the analog signal over the digital signal, the TMS-pulse Artifact appeared to start before the zero line, i.e., the time of the trigger. This is most likely due to the effect of a zero-phase filter which preserves the alignment of the peak while spreading the edges of the wave symmetrically on both sides. Therefore, increasing the sampling rate capacity of amplifiers could be useful for full characterization and subtraction of the TMS pulse.

Importantly, the reduction of the TMS-pulse Artifact below 5 ms revealed a short-duration Decay Artifact that was present even under optimal recording conditions. The initial amplitude of the Decay Artifact can reach the mV range, completely covering the EEG signal, which is a hundred times smaller.

The Decay Artifact can be reduced both with online and offline procedures. During recording, maintaining low impedances is crucial for shortening decays. Additionally, placing the wire direction perpendicular to the coil handle is helpful (Sekiguchi et al., 2011). However, these adjustments were implemented in this study and therefore appear insufficient to completely eliminate this artifact. The evidence from the data on tester that the decay artifact persisted when no gel was used suggests that it may depend on eddy currents within the electrode. Therefore, further developments in electrode material and shape may help reduce the Decay Artifact in future studies.

Regarding offline procedures (Hernandez-Pavon et al., 2022), the Decay Artifact is often removed with Independent Component Analysis (ICA) (Atluri et al., 2016; Rogasch et al., 2017, 2014; Wu et al., 2018). However, considering that the assumption of independence may be violated, ICA may not fully separate the artifact from physiological responses. Alternative approaches have been based on fitting procedures (Casula et al., 2017; Freche et al., 2018). Finally, the source-estimate-utilising-noise-discarding (SOUND) algorithm (Mutanen et al., 2018) can be employed to reduce the decay artifact if this is present in isolated electrodes. Crucially, the choice of the offline cleaning method can result in TEPs with different features (Bertazzoli et al., 2021; Casula et al., 2017; Hernandez-Pavon et al., 2012; Rogasch et al., 2022, 2014) and it is not clear yet which method may work better, although a first step to understand this has been recently published (Brancaccio et al., 2024).

Overall, more studies are needed to understand how online and/or offline procedures can effectively remove the Decay Artifact to recover cortical responses even earlier than the recently discovered immediate TMS-EEG responses (Beck et al., 2024a; Stango et al., 2024 preprint).

With this work we have added more information on the Early Artifacts, we were able to identify them and measure the overall duration. We hope that this could help scientists to better understand all the issues of the decay artifact and finally to find a good solution to eliminate it.

## Data availability statement

The data will be made available on an open repository on article acceptance.

## Author contributions

AS: Methodology, Formal analysis, Visualization, Writing - original draft

AZ: Conceptualization, Investigation, Formal analysis, Writing - original draft, Visualization

GB: Conceptualization, Investigation, Writing - original draft

SB: Formal analysis, Writing - review and editing

MB: Conceptualization, Data curation, Writing - original draft, Supervision, Project administration

