## Supplementary Material for "High-frequency sampling rate reduces TMS-pulse artifact duration but not decay artifact: implications for immediate TMS-EEG responses"

**Table S1.** Back to baseline (B2B) results. Main effects of Sampling rate and TMS intensity and their interaction, with post-hoc adjusted for multiple comparisons with Tuckey method.

|  | **Statistic** | ***p* value** |
| --- | --- | --- |
| **MELON** | | |
| **MagPro** |  |  |
| *Main effect Sampling rate* | *F = 20.8* | *p < 0.001* |
| *Post-hoc* |  |  |
| 4800 Hz vs. 9600 Hz | *t = -4.7* | *p < 0.001* |
| 4800 Hz vs. 19200 Hz | *t = 1.5* | *p = 0.31* |
| 9600 Hz vs. 19200 Hz | *t = 6.2* | *p < 0.001* |
| *Main effect TMS Intensity* | *F = 20.8* | *p < 0.001* |
| *Post-hoc* |  |  |
| 40 % vs. 70 % | *t = -4.8* | *p < 0.001* |
| 40 % vs. 100 % | *t = -6.2* | *p < 0.001* |
| 70 % vs. 100 % | *t = -1.4* | *p = 0.353* |
| *Interaction* | *F = 0.2* | *p = 0.957* |
| **Magstim** |  |  |
| *Main effect Sampling rate* | *F = 4.6* | *p = 0.011* |
| *Post-hoc* |  |  |
| 4800 Hz vs. 9600 Hz | *t = -1.7* | *p= 0.185* |
| 4800 Hz vs. 19200 Hz | *t = 1.3* | *p = 0.426* |
| 9600 Hz vs. 19200 Hz | *t = 3.0* | *p = 0.007* |
| *Main effect TMS Intensity* | *F = 20.9* | *p < 0.001* |
| *Post-hoc* |  |  |
| 40 % vs. 70 % | *t = -1.6* | *p = 0.23* |
| 40 % vs. 100 % | *t = -6.2* | *p < 0.001* |
| 70 % vs. 100 % | *t = -4.6* | *p < 0.001* |
| *Interaction* | *F = 4.7* | *p = 0.001* |
| *Post-hoc* |  |  |
| 19200 Hz: 100 % vs. 40 %  19200 Hz: 100% vs. 70 %  100%: 19200 Hz vs. 4800 Hz  100%: 19200 Hz vs. 9600 Hz  19200 Hz: 40% vs. 70%  40%: 19200 Hz vs. 4800 Hz  40%: 19200 Hz vs. 9600 Hz  70%: 9600 Hz vs. 4800 Hz  70%: 19200 Hz vs. 9600 Hz  4800 Hz: 100% vs. 40%  4800 Hz: 100% vs. 70 %  100%: 4800 Hz vs. 9600 Hz  4800 Hz: 40 % vs. 70%  40%: 4800 Hz vs. 9600 Hz  70%: 4800 Hz vs. 9600 Hz  9600 Hz: 100% vs. 40%  9600 Hz: 100% vs. 70%  9600 Hz: 40 % vs. 70% | *t = 1.1*  *t = -0.7*  *t = -3.6*  *t = -4.6*  *t = -1.8*  *t = 0.2*  *t = -0.7*  *t = 1.8*  *t = 0.0*  *t = 4.9*  *t = 4.7*  *t = -1.0*  *t = -0.2*  *t = -0.9*  *t = -1.8*  *t = 4.9*  *t = 3.9*  *t = -1.0* | *p = 0.974*  *p = 0.999*  *p = 0.012*  *p < 0.001*  *p = 0.711*  *p = 1.000*  *p = 0.998*  *p = 0.696*  *p = 1.000*  *p < 0.001*  *p < 0.001*  *p = 0.986*  *p = 1.000*  *p = 0.990*  *p = 0.709*  *p < 0.001*  *p = 0.003*  *p = 0.986* |
| **TESTER** | | |
| **MagPro** |  |  |
| *Main effect Sampling rate* | *F = 0.66* | *p = 0.515* |
| *Main effect TMS Intensity* | *F = 42.6* | *p < 0.001* |
| *Post-hoc* |  |  |
| 40 % vs. 70 % | *t = -5.5* | *p < 0.001* |
| 40 % vs. 100 % | *t =-9.2* | *p < 0.001* |
| 70 % vs. 100 % | *t = -3.7* | *p < 0.001* |
| *Interaction* | *F = 2.1* | *p = 0.079* |
| **Magstim** |  |  |
| *Main effect Sampling rate* | *F = 3.4* | *p = 0.036* |
| *Post-hoc* |  |  |
| 4800 Hz vs. 9600 Hz | *t = 0.3* | *p= 0.955* |
| 4800 Hz vs. 19200 Hz | *t = -2.1* | *p = 0.097* |
| 9600 Hz vs. 19200 Hz | *t = -2.4* | *p = 0.045* |
| *Main effect TMS Intensity* | *F = 88.0* | *p < 0.001* |
| *Post-hoc* |  |  |
| 40 % vs. 70 % | *t = -3.3* | *p = 0.003* |
| 40 % vs. 100 % | *t = -10.1* | *p < 0.001* |
| 70 % vs. 100 % | *t = -6.9* | *p < 0.001* |
| *Interaction* |  |  |
| *Post-hoc* |  |  |
| 19200 Hz: 100 % vs. 40 %  19200 Hz: 100% vs. 70 %  100%: 19200 Hz vs. 4800 Hz  100%: 19200 Hz vs. 9600 Hz  19200 Hz: 40% vs. 70%  40%: 19200 Hz vs. 4800 Hz  40%: 19200 Hz vs. 9600 Hz  70%: 9600 Hz vs. 4800 Hz  70%: 19200 Hz vs. 9600 Hz  4800 Hz: 100% vs. 40%  4800 Hz: 100% vs. 70 %  100%: 4800 Hz vs. 9600 Hz  4800 Hz: 40 % vs. 70%  40%: 4800 Hz vs. 9600 Hz  70%: 4800 Hz vs. 9600 Hz  9600 Hz: 100% vs. 40%  9600 Hz: 100% vs. 70%  9600 Hz: 40 % vs. 70% | *t = 8.7*  *t = 6.4*  *t = 3.1*  *t = 5.3*  *t = -2.3*  *t = 0.5*  *t = -0.6*  *t = 0.0*  *t = -0.4*  *t = 6.2*  *t = 3.4*  *t = 2.2*  *t = -2.7*  *t = -1.1*  *t = -0.5*  *t = 2.8*  *t = 0.7*  *t = -2.1* | *p < 0.001*  *p < 0.001*  *p = 0.050*  *p < 0.001*  *p = 0.342*  *p = 1.000*  *p = 0.999*  *p = 1.000*  *p = 1.000*  *p < 0.001*  *p = 0.019*  *p = 0.403*  *p = 0.138*  *p = 0.972*  *p = 1.000*  *p = 0.106*  *p = 0.998*  *p = 0.475* |

**Table S2.** End of TMS-pulse Artifact (ETA) results. Main effects of Sampling rate and TMS intensity and their interaction, with post-hoc adjusted for multiple comparisons with Tuckey method.

|  | **Statistic** | **p value** |
| --- | --- | --- |
| **MELON** | | |
| **MagPro** |  |  |
| *Main effect Sampling rate* | *F = 558.4* | *p < 0.001* |
| *Post-hoc* |  |  |
| 4800 Hz vs. 9600 Hz | *t = 16.7* | *p < 0.001* |
| 4800 Hz vs. 19200 Hz | *t = 33.5* | *p < 0.001* |
| 9600 Hz vs. 19200 Hz | *t = 16.8* | *p < 0.001* |
| *Main effect TMS Intensity* | *F = 77.0* | *p < 0.001* |
| *Post-hoc* |  |  |
| 40 % vs. 70 % | *t = 10.0* | *p < 0.001* |
| 40 % vs. 100 % | *t = 11.4* | *p < 0.001* |
| 70 % vs. 100 % | *t = 1.3* | *p* = *0.386* |
| *Interaction* | *F = 43.7* | *p < 0.001* |
| *Post-hoc* |  |  |
| 19200 Hz: 100 % vs. 40 %  19200 Hz: 100% vs. 70 %  100%: 19200 Hz vs. 4800 Hz  100%: 19200 Hz vs. 9600 Hz  19200 Hz: 40% vs. 70%  40%: 19200 Hz vs. 4800 Hz  40%: 19200 Hz vs. 9600 Hz  70%: 9600 Hz vs. 4800 Hz  70%: 19200 Hz vs. 9600 Hz  4800 Hz: 100% vs. 40%  4800 Hz: 100% vs. 70 %  100%: 4800 Hz vs. 9600 Hz  4800 Hz: 40 % vs. 70%  40%: 4800 Hz vs. 9600 Hz  70%: 4800 Hz vs. 9600 Hz  9600 Hz: 100% vs. 40%  9600 Hz: 100% vs. 70%  9600 Hz: 40 % vs. 70% | *t = -15.5*  *t = 0.3*  *t = -23.6*  *t = -11.9*  *t = 15.8*  *t = -11.3*  *t = 2.3*  *t = -26.9*  *t = -11.6*  *t = -3.2*  *t = -3.0*  *t = 11.7*  *t = 0.2*  *t = 13.6*  *t = 15.4*  *t = -1.4*  *t = 0.6*  *t = 2.0* | *p < 0.001*  *p* = *1.000*  *p < 0.001*  *p < 0.001*  *p < 0.001*  *p < 0.001*  *p* = *0.372*  *p < 0.001*  *p < 0.001*  *p* = *0.041*  *p* = *0.067*  *p < 0.001*  *p* = *1.000*  *p < 0.001*  *p < 0.001*  *p* = *0.913*  *p* = *0.999*  *p* = *0.555* |
| **Magstim** |  |  |
| *Main effect Sampling rate* | *F = 331.6* | *p < 0.001* |
| *Post-hoc* |  |  |
| 4800 Hz vs. 9600 Hz | *t = 17.5* | *p < 0.001* |
| 4800 Hz vs. 19200 Hz | *t = 25.1* | *p < 0.001* |
| 9600 Hz vs. 19200 Hz | *t = 7.6* | *p < 0.001* |
| *Main effect TMS Intensity* | *F = 7.0* | *p < 0.001* |
| *Post-hoc* |  |  |
| 40 % vs. 70 % | *t = 2.5* | *p = 0.032* |
| 40 % vs. 100 % | *t = -1.1* | *p = 0.486* |
| 70 % vs. 100 % | *t = -3.7* | *p < 0.001* |
| *Interaction* | *F = 6.0* | *p < 0.001* |
| *Post-hoc* |  |  |
| 19200 Hz: 100 % vs. 40 %  19200 Hz: 100% vs. 70 %  100%: 19200 Hz vs. 4800 Hz  100%: 19200 Hz vs. 9600 Hz  19200 Hz: 40% vs. 70%  40%: 19200 Hz vs. 4800 Hz  40%: 19200 Hz vs. 9600 Hz  70%: 9600 Hz vs. 4800 Hz  70%: 19200 Hz vs. 9600 Hz  4800 Hz: 100% vs. 40%  4800 Hz: 100% vs. 70 %  100%: 4800 Hz vs. 9600 Hz  4800 Hz: 40 % vs. 70%  40%: 4800 Hz vs. 9600 Hz  70%: 4800 Hz vs. 9600 Hz  9600 Hz: 100% vs. 40%  9600 Hz: 100% vs. 70%  9600 Hz: 40 % vs. 70% | *t = 0.6*  *t = 2.6*  *t = -14.3*  *t = -2.7*  *t = 2.0*  *t = -14.2*  *t = -4.7*  *t = -16.7*  *t = -6.8*  *t = 0.7*  *t = 0.2*  *t = 11.6*  *t = -0.5*  *t = 9.6*  *t = 9.9*  *t = -1.3*  *t = -1.5*  *t = -0.2* | *p = 0.999*  *p = 0.173*  *p < 0.001*  *p = 0.137*  *p = 0.545*  *p < 0.001*  *p < 0.001*  *p < 0.001*  *p < 0.001*  *p = 0.999*  *p = 1.000*  *p < 0.001*  *p = 1.000*  *p < 0.001*  *p < 0.001*  *p = 0.936*  *p = 0.875*  *p = 1.000* |
| **TESTER** | | |
| **MagPro** |  |  |
| *Main effect Sampling rate* | *F = 885.8* | *p < 0.001* |
| *Post-hoc* |  |  |
| 4800 Hz vs. 9600 Hz | *t = 22.5* | *p < 0.001* |
| 4800 Hz vs. 19200 Hz | *t = 42.1* | *p < 0.001* |
| 9600 Hz vs. 19200 Hz | *t = 19.6* | *p < 0.001* |
| *Main effect TMS Intensity* | *F = 6.1* | *p = 0.002* |
| *Post-hoc* |  |  |
| 40 % vs. 70 % | *t = -3.27* | *p = 0.003* |
| 40 % vs. 100 % | *t = -2.7* | *p = 0.019* |
| 70 % vs. 100 % | *t = 0.6* | *p = 0.837* |
| *Interaction* | *F = 0.6* | *p = 0.678* |
| **Magstim** |  |  |
| *Main effect Sampling rate* | *F =243.9* | *p < 0.001* |
| *Post-hoc* |  |  |
| 4800 Hz vs. 9600 Hz | *t = 18.2* | *p < 0.001* |
| 4800 Hz vs. 19200 Hz | *t = 19.9* | *p < 0.001* |
| 9600 Hz vs. 19200 Hz | *t = 1.8* | *p = 0.168* |
| *Main effect TMS Intensity* | *F = 39.6* | *p < 0.001* |
| *Post-hoc* |  |  |
| 40 % vs. 70 % | *t = -0.8* | *p = 0.726* |
| 40 % vs. 100 % | *t = -8.1* | *p < 0.001* |
| 70 % vs. 100 % | *t = -7.3* | *p < 0.001* |
| *Interaction* | *F=13.1* | *p < 0.001* |
| *Post-hoc* |  |  |
| 19200 Hz: 100 % vs. 40 %  19200 Hz: 100% vs. 70 %  100%: 19200 Hz vs. 4800 Hz  100%: 19200 Hz vs. 9600 Hz  19200 Hz: 40% vs. 70%  40%: 19200 Hz vs. 4800 Hz  40%: 19200 Hz vs. 9600 Hz  70%: 9600 Hz vs. 4800 Hz  70%: 19200 Hz vs. 9600 Hz  4800 Hz: 100% vs. 40%  4800 Hz: 100% vs. 70 %  100%: 4800 Hz vs. 9600 Hz  4800 Hz: 40 % vs. 70%  40%: 4800 Hz vs. 9600 Hz  70%: 4800 Hz vs. 9600 Hz  9600 Hz: 100% vs. 40%  9600 Hz: 100% vs. 70%  9600 Hz: 40 % vs. 70% | *t = 12.2*  *t = 10.1*  *t = -5.7*  *t = 3.5*  *t = -2.2*  *t = -17.2*  *t = -5.4*  *t = -14.9*  *t = -4.0*  *t = 1.2*  *t = 1.3*  *t = 9.5*  *t = 0.0*  *t = 11.7*  *t = 11.0*  *t = 3.5*  *t = 2.7*  *t = -0.7* | *p < 0.001*  *p < 0.001*  *p < 0.001*  *p = 0.014*  *p = 0.425*  *p < 0.001*  *p < 0.001*  *p < 0.001*  *p = 0.003*  *p = 0.948*  *p = 0.936*  *p < 0.001*  *p = 1.000*  *p < 0.001*  *p < 0.001*  *p = 0.017*  *p = 0.134*  *p = 0.999* |
